# Generation of fluorescent cell-derived-matrix to study 3D cell migration

**DOI:** 10.1101/746941

**Authors:** Amélie Luise Godeau, Hélène Delanoë-Ayari, Daniel Riveline

## Abstract

Cell migration is involved in key phenomena in biology, ranging from development to cancer. Fibroblasts move between organs in 3D polymeric networks. So far, motile cells were mainly tracked *in vitro* on Petri dishes or on coverslips, *i.e.* 2D flat surfaces, which made the extrapolation to 3D physiological environments difficult. We therefore prepared 3D Cell Derived Matrix (CDM) with specific characteristics with the goal of extracting the main readouts required to measure and characterise cell motion: cell specific matrix deformation through the tracking of fluorescent fibronectin within CDM, focal contacts as the cell anchor and acto-myosin cytoskeleton which applies cellular forces. We report our method for generating this assay of physiological-like gel with relevant readouts together with its potential impact in explaining cell motility *in vivo*.

## 1 Introduction

### 1.1 Cell migration: 2D versus 3D

Cell migration has been characterised on Petri dishes and 2D flat surfaces for more than a century (Grote, 2018; Petri, 1887). In this condition, cells are easily tracked within single focal planes with phase contrast microscopy. Their culture is accessible to any laboratory with standard equipment and since the seminal studies of Michael Abercrombie (Abercrombie & Heaysman, 1953), this test remains the classical and ‘easy’ assay for the study of cell motility.

However, cells evolve in 3D environments *in vivo, i.e*. usually in dense polymeric networks. This change in geometry and dimensions implies numerous differences between motions in 2D and in 3D conditions: (i) distributions of adhesion zones - focal contacts - are located at the cell-substrate interface in 2D and all around the cell surface in 3D; (ii) compositions of adhesion contacts can be distinct; (iii) 3D networks entail physical constraints for cells (Charras & Sahai, 2014) which need to generate their own space by potentially squeezing nuclei (Wolf et al., 2013) or by cleaving their environments with metalloproteinase (Nagase, Visse, & Murphy, 2006); (iv) 2D flat surfaces are very rigid compared to soft physiological matrices, and this can change motion and the associated dynamics of the cytoskeleton.

These differences come along with distinct cellular structures responsible for cell motility. In the classical view on 2D surfaces, cells adhere, they generate lamellipodia at their front which extend through actin polymerisation, myosin II based contraction contributes to deform the cell body forward, and focal adhesions at the back of the cell are disassembled (Ridley et al., 2003). For 3D environments, the picture is more complex and cells show multiple strategies to move. Indeed, cell migration in 3D has been studied in different configurations: in gels (Lämmermann et al., 2008; Petrie, Gavara, Chadwick, & Yamada, 2012; Wolf et al., 2013), in micro-channels (Lautenschlager et al., 2009; Liu et al., 2015; Prentice-Mott et al., 2013), or cells ‘sandwiched’ between two plates (Yip, Chiam, & Matsudaira, 2015). In these assays, various types of migration were reported, such as mesenchymal migration, amoeboid migration (Liu et al., 2015), or bleb-based (Petrie et al., 2012) motion. Each type of migration involves distinct distributions of forces applied by cells (see for example (Bergert et al., 2015; Liu et al., 2015)). This variety of behaviours calls for new assays to connect cellular activity, local force application and cell motion.

During movement, cells apply local forces, which can be decomposed in moments (force monopole, dipole, quadrupole) to generate a motion with zero total force, as set by laws of motion at low Reynolds number (Tanimoto & Sano, 2014). To identify the composition of these forces, traction force microscopy allows to capture deformations in the local cellular environment. Physical descriptions have emerged for explaining motion in connection with force patterns or cellular structures (Saez, Buguin, Silberzan, & Ladoux, 2005; Tanimoto & Sano, 2014).

Along these lines, we decided to design an assay where essential readouts would be acquired and integrated, *i.e*. matrix deformation and cellular actors generating motility, to understand cell migration. We adapted a CDM protocol (Cukierman, Pankov, Stevens, & Yamada, 2001) where cells move within a thin layer of physiological matrix in order to visualize the main anchor protein for cells in the extracellular matrix, *i.e*. fibronectin (FN). We plated cells and observed their motion and the associated deformation of the FN network over hours and days. Altogether, this assay captures spatio-temporal evolution of deformation, anchoring points in the extracellular matrix (ECM), *focal contact* and acto-myosin distributions. This approach could serve as a generic method to study and characterise cell motion in 3D environments *in vitro*.

### 1.2 The need for *in vitro* assays

In order to understand cell motility in 3D, there is a necessity to design specific culture set-ups allowing to control the mechanical and biochemical environments. The mechanical state can be tuned by adjusting Young modulus of the matrix, and the biochemical composition and state can be modulated by controlling synthesis and growth conditions. To study the implication of forces on cell motility, a substrate or network which can be deformed by cells is required. Considering that cells apply pN forces (Balaban et al., 2001; Riveline et al., 2001), the gel should have a low and known Young’s Modulus in order to detect forces applied by cells. Cell derived matrices (CDM) have a considerable potential for cell studies in 3D physiological environment (Cukierman et al., 2001). These matrices are produced by cells, which are removed after days thereby leaving a 3D-like physiological gel. This set-up has already been used for studying focal adhesion distributions in 3D (Cukierman et al., 2001). Designing a cellular environment with low Young’s modulus and a visualisation of the relevant extracellular matrix protein – fibronectin - allows to address specific questions regarding symmetry breaking, spatio-temporal force patterns and cell adhesion in 3D. We achieved this goal by adapting cell derived matrix protocols, which have been shown to generate low Young’s modulus with suitable size to embed cells within CDM while allowing migration along x-y plane fitting the cell body.

We prepared and visualised this fibronectin network in two ways. We introduced fluorescently labelled fibronectin to the matrix culture, and we generated a genetically modified cell line which produces and secretes fluorescently labelled fibronectin. We report these methods in this article. In the next section (section 2), we introduce the general matrix protocol, and we explain how to visualise the fluorescent fibronectin within these matrices. This is followed by the description of introduction of beads in the CDM for its mechanical characterisation. In section 3, we explain how to integrate cells in our CDM, and how to set up a time-lapse movie. We also give an example of how to process the data via the Kanade-Lucas-Tomasi (KLT) feature tracker to extract matrix deformation. In section 4, we present and discuss some of our results, and in section 5, we give an outlook of potential applications.

## 2 Generation of fluorescent CDM

In the following section, we first explain the method for culturing CDM (Cukierman et al., 2001, Castelló-Cros & Cukierman, 2009; Goetz et al., 2011). We then describe (i) how to modify this protocol in order to obtain fluorescent matrices; and (ii) how to integrate micron-sized beads within the matrix to generate an external framework for correcting the sample drift or for mechanical characterisation.

### 2.1 Preparation of CDM

#### 2.1.1 Material

○ 6x multiwell plates or 20mm Petri dishes
○ Pipettes
○ Glass coverslip
○ Gelatin 1%
○ Glutaraldehyde 1%
○ Glycine
○ PBS 1%
○ NIH3T3 fibroblasts
○ L-ascorbic acid
○ DMEM
○ Bovine calf serum (BCS)
○ Penicillin and streptomycin (pen-strep)
○ Trypsin-EDTA
○ Triton
○ NH_4_OH

#### 2.1.2 Equipment

○ Cell incubator
○ Laminar flow hood
○ Centrifuge
○ Fridge 4°C

#### 2.1.3 Method: Matrix culture

These steps are conducted in sterile conditions under a laminar flow hood. All reagents have been autoclaved or filtered before usage to prevent contamination. In this protocol, we take advantage that fibroblasts produce and secrete extracellular matrix proteins. These proteins are then organized into a physiological-like matrix by cells. To keep and stabilise this matrix, cells are cultured on a gelatin-based support. The protocol is schematically represented in Figure 1: seeding of cells on the gelatin-based substrate in order to obtain a confluent monolayer, culture of CDM and cells removal.

1. For microscopy, we use glass coverslips (CS) as a substrate to ease experiments with high resolution. Multiwell plates or Petri dishes can be used as well. A glass coverslip (#1) is covered with 1ml of 1% gelatin (gelatin from cold water fish skin, Sigma) and placed at 37°C in the incubator for 1 h to promote anchorage of the future CDM. After the CS has been washed twice with 1% PBS, the gelatin is incubated for 20 min at room temperature with 1% glutaraldehyde (Sigma). This chemical agent crosslinks gelatin and provides stability to the scaffold. The CS is rinsed again twice with PBS before incubation for 20-30 min with 1 M glycine (Sigma) at room temperature. This allows to saturate the aldehyde groups generated by the glutaraldehyde treatment in gelatin. Subsequently the CS is washed twice with PBS. At this step, the protocol can be continued, or alternatively the CS can be stored at 4°C up to one month.
2. A Petri dish with an 80% confluent monolayer is taken and washed 1x with PBS. PBS is removed and trypsin-EDTA solution is added. The dish is incubated for 3 min at 37°C in the incubator to detach cells from the surface. The supernatant is recovered and added to a 15ml tube with DMEM supplemented with 10% BCS and 1% penicillin-streptomycin solution. Cells suspended in cell culture medium are centrifuged for 5 min, the supernatant removed and the cell pellet resuspended in DMEM supplemented with 10% BCS and 1% pen-strep. Cells are plated on the gelatin-coated CS. They need to be plated at high density in order to produce CDM. For NIH3T3 cells, this corresponds to 1 million cells for 35 x 10 mm Petri dish.
3. This sample is placed in the incubator at 37°C to let cells adhere. Every second day, the culture medium supplemented with 50 *μ*g/ml L-ascorbic acid is changed. L-ascorbic acid is added to promote collagen secretion of cells. The culture is maintained for 8-9 days. During this time, the synthesis and accumulation of CDM can be observed.
4. Cells are removed by lysis with a medium consisting of 20 mM NH_4_OH and 0.5 % Triton (Sigma) in 1x PBS after washing twice with PBS. NH_4_OH stabilises CDM in the presence of Triton. The prewarmed lysis medium is carefully pipetted on the CS and incubated for up to 10 min at 37°C in the incubator. The cell lysis (Figure 2 and Movie 1) is rapid, and the extrusion of nuclei can be observed. If the CDM culture (step 3) has been maintained for too long, the CDM might become too dense and nuclei might stay trapped within the CDM. PBS is added and the CDM stored overnight at 4°C: this ensures complete lysis of cellular membranes and provides more temporal stability to the CDM. The day after, PBS is carefully changed 3 times to remove all residues of Triton to avoid undesired effects on cells during experiments. Matrices are covered with PBS and kept at 4°C. Although CDM can be stored for up to 1 month, it is recommended to use them in the following two weeks, as risks of detachment of the matrix from the CS increase with time.

**Figure 1:**
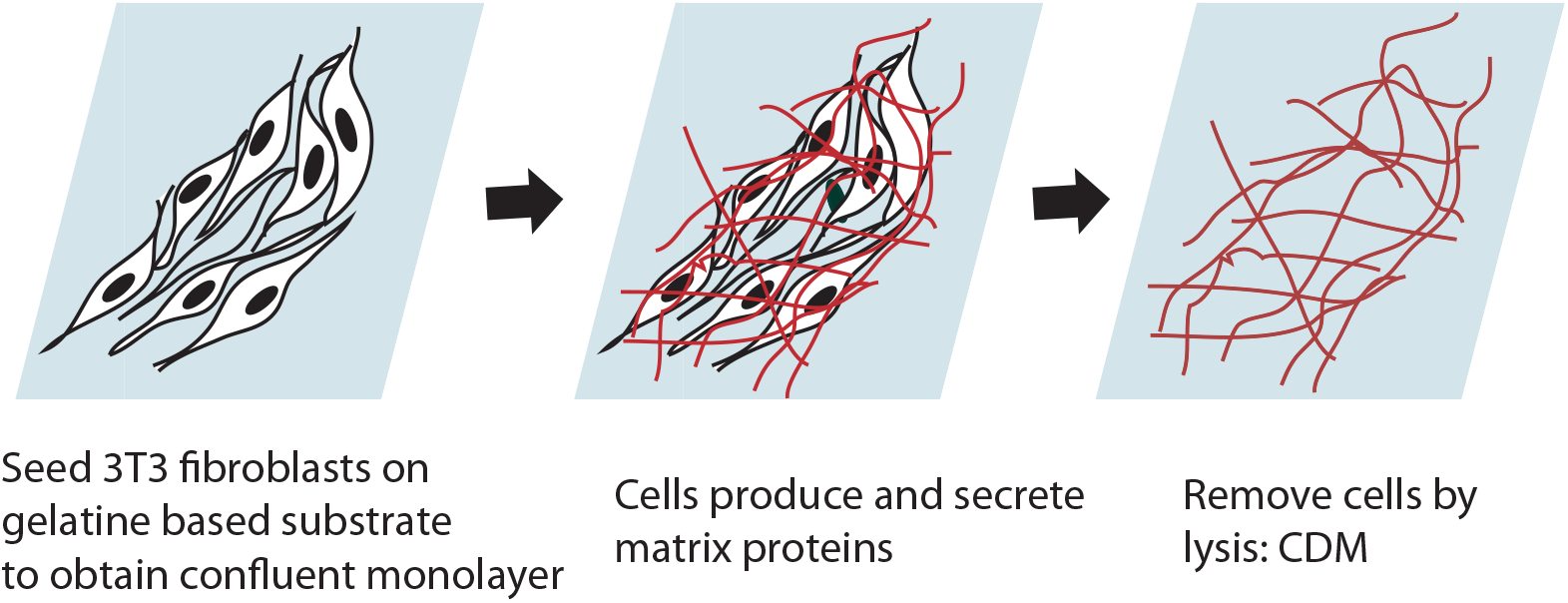
Schematic description of CDM production. The glass coverslip is covered with gelatin and further treatments with glutaraldehyde and glycine are performed before cells are seeded. Culture is maintained 8-9 days before cells lysis, leaving only the secreted proteins forming the CDM.

**Figure 2:**
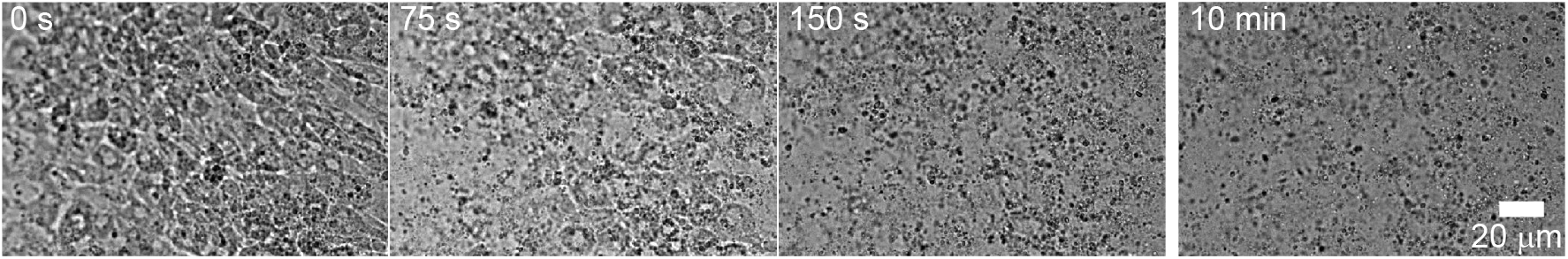
Lysis of cells in CDM: images of cells lysis producing CDM. Most of the membranes are lysed within the first 2-3 min of the process and after 10 min no cell membranes or nuclei are visible anymore; see also Movie 1.

### 2.2 Generation of a fluorescent matrix

In the following section, we present two methods to generate cell derived matrices with fluorescently labelled fibronectin. Their quality can be evaluated with immunofluorescence microscopy by comparing the spatial localisations of fibronectin with both labels (data not shown). We add a rhodamine labelled fibronectin to the culture medium (Figure 3a) or we modify genetically cells for making them secrete fluorescent fibronectin (Figure 3b).

**Figure 3:**
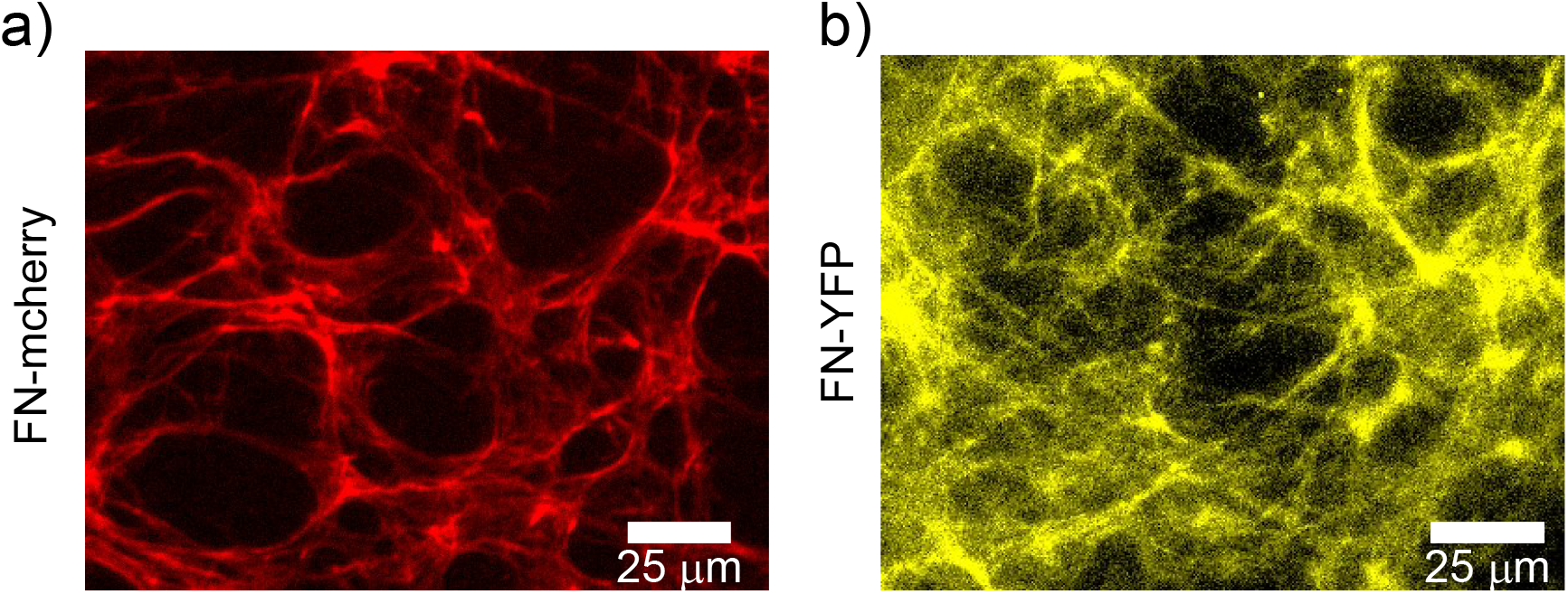
Fluorescent fibronectin within CDMs. a) Image of fibronectin mesh acquired with a x20 objective. Fluorescent fibronectin added to culture medium is integrated in matrix. b) Image of fibronectin mesh acquired with a x20 objective secreted by genetically modified cells. These cells synthesise and secrete fluorescent fibronectin which is incorporated in the matrix.

#### 2.2.1 Material

○ Rhodamine labeled fibronectin (TRITC fibronectin, Cytoskeleton)
○ Fluorescent fibronectin expressing NIH3T3 cells
○ And the material of section 2.1.1

#### 2.2.2 Equipment

○ Cell incubator
○ Laminar flow hood
○ Centrifuge
○ Fridge 4°C

#### 2.2.3 Method

1. For this first method, step 1 and 2 are performed as described in section 2.1.3. Step 3 is modified as follows: The culture is scaled down to fit a 96 well-plate. Normal NIH3T3 fibroblasts are used for CDM production. Additionally, to the L-Ascorbic acid the medium is supplemented with rhodamine labelled fibronectin (1 *μ*g per 100 μl well) for every change of medium. Cells producing CDM incorporate to the matrix the fluorescent fibronectin present in the medium. After 8-9 days of culture, we proceed to step 4 as described above. When storing at 4°C, the Petri dishes or multiwell plates containing the CDM are surrounded with aluminum foil to prevent photo-bleaching.
2. The second method consists in using genetically-modified cells which produce and secrete fluorescently labeled fibronectin. Protocols for establishing stable cell lines can be found elsewhere (Ruedel & Bosserhoff, 2012). Briefly, we transfected NIH3T3 fibroblasts with a fluorescent Ypet-FN construct from Erickson laboratory (Ohashi, Kiehart, & Erickson, 1999) using Lipofectamin (Invitrogen Gibco, Thermo Fisher Scientific). The method follows the steps in section 2.1.3, but cells stably expressing FN-Ypet are seeded onto the CS for the CDM production in step 3, instead of using regular NIH3T3 fibroblasts. To select cells encoding the Ypet-FN, cell culture media contain G418 antibiotics. After cloning, this stable cell line is used for the preparation of CDM. The fluorescently labelled fibronectin is then secreted and spontaneously incorporated the CDM. When the matrix has been produced, cells are removed as described in step 4 of section 2.1.3. During storage, we surround the Petri dishes or multiwell plates containing the CDM with aluminum foil to prevent photo-bleaching.

### 2.3 Insertion of beads in the CDM

Beads within matrices can be used for different purposes. They can serve as markers for calculating the spatial drift potentially occurring during time-lapse experiments. They can also be used as traps for optical or magnetic tweezers for mechanical characterisation of CDM, or also as fiducial markers to calculate the matrix displacement generated by external forces. We recommend to use monochromatic beads to make sure that there is no colour overlap between fluorescent channels and facilitate further analysis.

#### 2.3.1 Material

○ Fluorescent or magnetic beads of typical 1μm diameter or below

#### 2.3.2 Equipment

○ Plasma cleaner
○ Spin-coater
○ Oven

#### 2.3.3 Method

1. For drift correction, beads need to stick to the surface of coverslips and not to the matrix, nor to be loosely trapped within the matrix. The coverslips need to be activated by plasma cleaning to make the glass hydrophilic. A bead solution is prepared with a dilution of 1:1000 (to be adjusted with bead size). The CS is placed on a spin-coater and a drop of approximately 200μl of the bead solution is spin-coated at 1000 rpm for 1 min on the CS, and the sample is dried in an oven at 65°C for at least 30 min. Then CS with beads are exposed to UV light for at least 15 min to prevent contamination during CDM culture. Since bead fluorescent intensity is high and stable, the risk of photo-bleaching is low. Afterwards, the protocol from section 2.1.3 can be followed.
2. There is also the option to incorporate fluorescent or magnetic beads in the matrix. Therefore, here again step 3 of the protocol in section 2.1.3 is modified. Fluorescently labelled beads (or any other beads) are added to the culture medium. As described before for fluorescently labeled fibronectin, beads integrate the CDM while cells secrete the matrix proteins. Beads are incorporated throughout the height/thickness of the CDM. It may happen that incorporated beads are not well anchored to the matrix and loosely diffuse within the meshwork. Fluorescent beads can also be used to calculate displacements applied to the matrix and serve as proxys for the matrix/fiber localization. They can as well be employed for mechanical characterisation of the matrix such as magnetic or optical tweezer measurements.

## 3 Characterisation of cell migration in fluorescent CDM

When the fluorescent CDM is produced, cells can be incorporated within the matrix. Their motility and force application can then be studied. In the following section, we explain how to integrate cells in the CDM, how to track their migration and the matrix deformation under the microscope, and how to process the data in order to extract the matrix displacement. It is highly recommended to use cells which have been transfected with a construct leading to the expression of relevant fluorescent proteins: we transfected for proteins of the cytoskeleton (actin, microtubule), motor proteins (myosin) or proteins revealing focal adhesions (zyxin, see Figure 5)). This allows to visualise not only the fibronectin of the matrix but also cellular structures which generate forces on the CDM and to follow quantitatively the spatio-temporal behavior of these physical readouts.

### 3.1 Integration of cells in the CDM

Cell medium and trypsin are heated in a water bath to 37°C and CDMs placed at room temperature for at least 1h. The microscope is pre-heated to 37°C for at least 1h (and if possible overnight) to reach stable imaging conditions and to minimise drift during the acquisition.

#### 3.1.1 Material

○ NIH3T3 fibroblasts (or any other cell types – see Figure 4)
○ Leibowitz 15 (L-15) medium
○ Bovine calf serum (BCS)
○ Trypsin-EDTA
○ Penicillin streptomycin (Pen-Strep)

**Figure 4:**
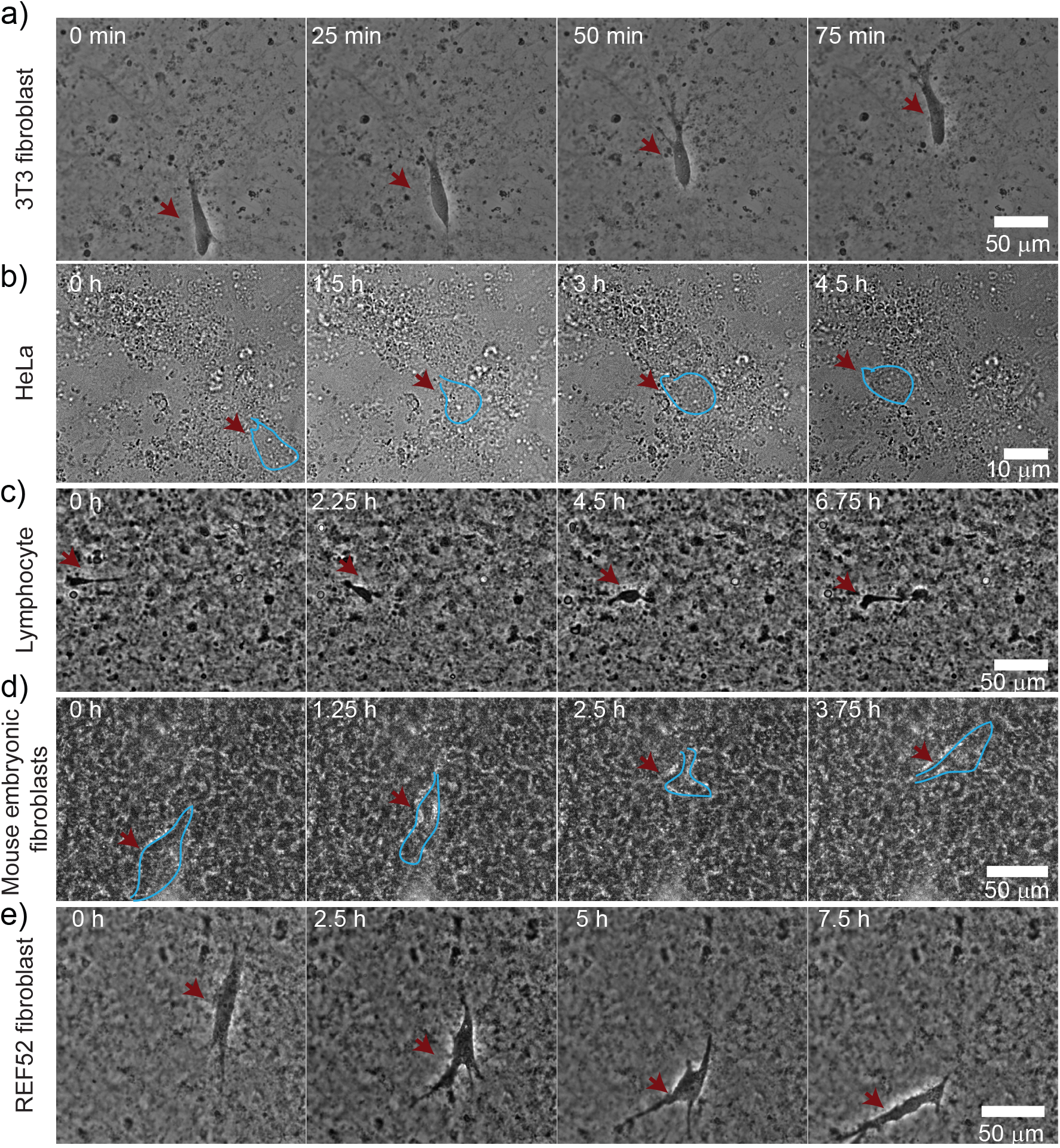
Different cell types migrate in CDM: a) NIH3T3, b) HeLa, c) mouse lymphocytes, d) mouse embryonic fibroblasts and e) REF52 fibroblast integrate and show motility in our CDM, see also Movies 2-6. All cell positions are indicated with a red arrow some cells are outlined in light blue.

#### 3.1.2 Equipment

○ Cell incubator
○ Laminar flow hood
○ Centrifuge
○ Metallic holder
○ Microscope

#### 3.1.3 Method

We prepare cells in a Petri dish reaching 80% confluent monolayer. We then plate cells at 100 cells/mm^2^ density. PBS is removed, and the CDM is incubated with cells for at least 2h at 37°C. CS with CDM and cells is mounted on a metallic holder which has to be cleaned with ethanol in a sonicator, dried and then exposed to UV light to reduce possible contamination during experiment. If the microscope is used with CO_2_, the medium is changed to DMEM supplemented with 1% BCS and 1% penicillin streptomycin; otherwise, we used L-15 medium supplemented with 1% BCS and 1% pen-strep allowing for CO_2_-free experimental conditions. Subsequently CDM with cells is mounted on the microscope and time-lapse experiment is started. A variety of cells move within these CDMs, stable cell lines, primary cells, fibroblasts and epithelial cells (see Figure 4 and Movies 2-6). For the investigation of cell integration in CDM, cells are resuspended in L15 after trypsinisation and added to the CDM under the microscope. The rapid incorporation of cells within the matrix can be observed as shown in Figure 5 and Movie 7.

**Figure 5:**
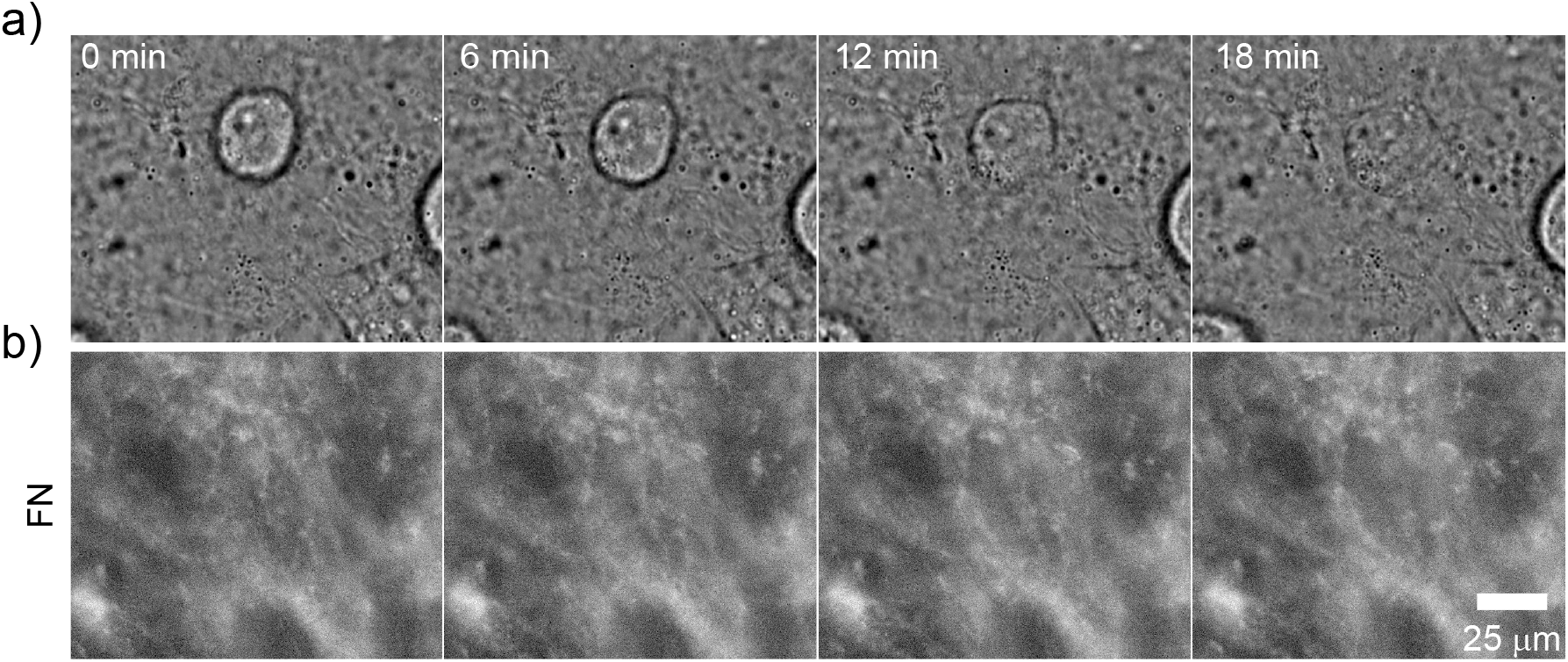
Cells enter the CDM. a) bright field image and b) fibronectin. At time zero, the cell is spherical on top of the CDM; entry occurs within minutes; see also Movie 7.

### 3.2 Cell migration within CDM

Cell migration is recorded via time-lapse experiments using microscopes. For high resolution and limitation of photo-bleaching, we recommend confocal microscopy and spinning disk microscope. To capture cell migration and deformation of the matrix, a z-stack spanning the CDM thickness is recorded. The frequency of image acquisition is adapted to experimental requirements. At least two channels are used during experiments to differentiate the matrix from cells. In order to have an optimal experiment, photo-bleaching, photo-toxicity and temporal requirements need to be balanced and frequency of image acquisition, exposure time and intensity need to be adapted. As an illustration, typical migration of a cell expressing RFP-zyxin in red in a CDM with fluorescent fibronectin in yellow is shown in Figure 6 and in Movie 8.

**Figure 6:**
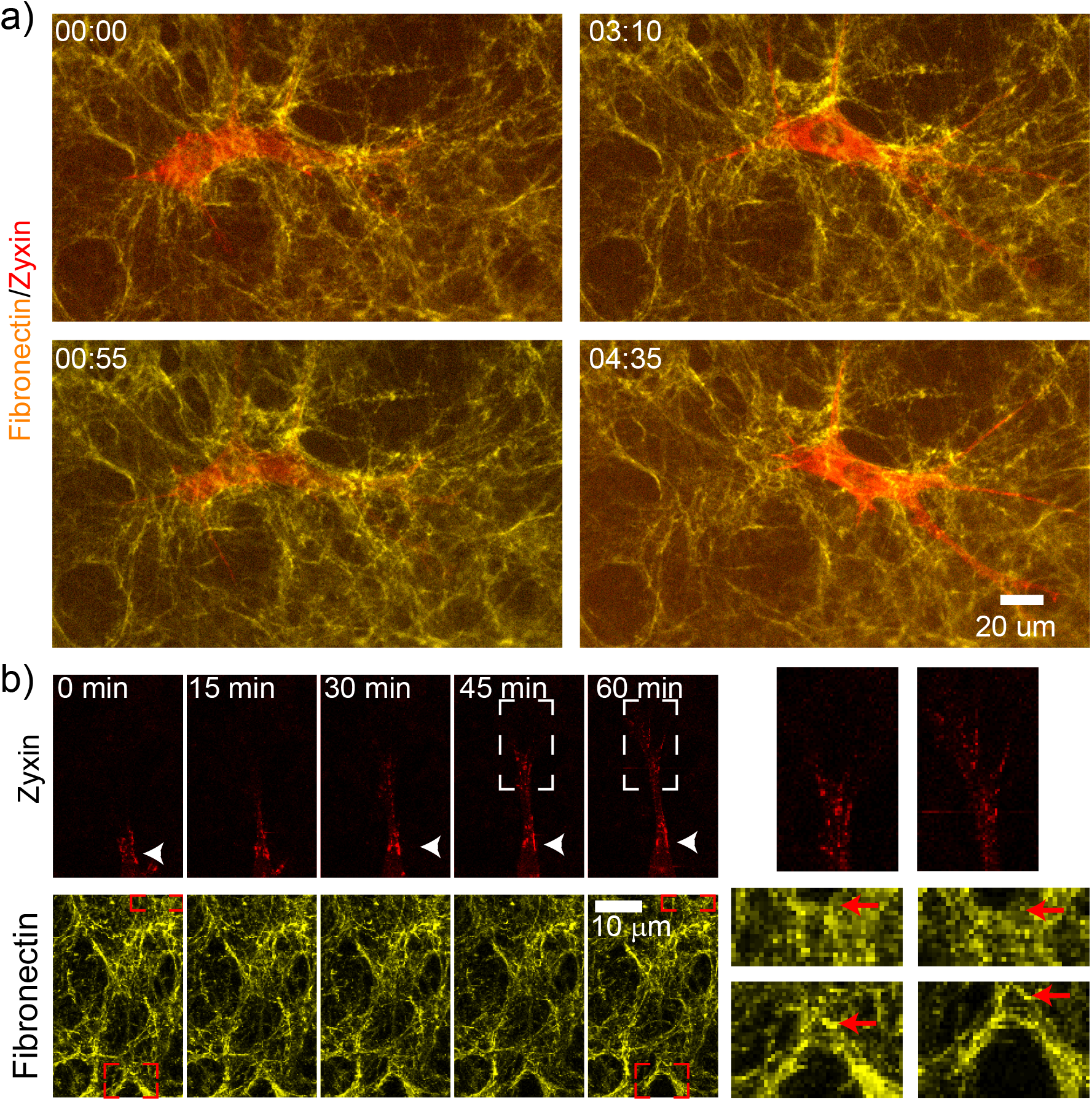
Cell migrating in CDM: a) Time-lapse images of a cell expressing RFP-zyxin a protein of focal adhesions in red, migrating in the fluorescent fibronectin in yellow embedded within the CDM. Time in hh:mm. b) Image sequence of focal adhesion dynamics (top) and growth highlighted in the zoomed region above white arrow for reference, and (bottom) corresponding matrix displacement during cell migration, with enlarged region (right, matrix displacement highlighted with red arrows); see also Movie 8.

### 3.3 Calculation of matrix displacement

To assess the deformation or the displacement of matrix, different methods can be used, such as Particle Imaging Velocimetry (PIV) (Pakshir et al., 2019; Steinwachs et al., 2015), other detection methods (Legant et al., 2013, 2010) or custom KLT feature tracker (Godeau et al., in preparation). Before these calculations, images are pre-treated (i) to generate images on which matrix displacements are then evaluated and (ii) to correct for any artifact due to the characteristics of the imaging system or the imaging conditions.

#### 3.3.1.1 Image pre-treatments

Before extracting the deformation or displacement of the matrix, the raw data of matrix needs to be either pre-treated or processed. The aim of the pre-treatment is to obtain 2D images for the calculation of x-y displacement of the mesh with the best possible signal-to-noise ratio. The pre-treatment options consider images acquired with multi-positions, z-stack and images with artifacts.

- If images have been obtained with multi-positions, stitching of the images needs to be performed to obtain a continuous image.
- If a z-stack has been acquired, the different planes need to be projected on one plane. This can be done using Fiji with the z-projection plugin with maximal or average projection.
- If artifacts like ‘comets’ or bright spots can be seen in images, they can be removed by choosing the “remove outlier” or “de-speckle” option in Fiji noise plugin.
- If the frame of the image is very large compared to the area of deformation, it can be resized to reduce calculation load of the next step.

#### 3.3.1.2 Tracker algorithm

In the following, we focus on the KLT tracker method as PIV methods have already been discussed elsewhere (Pakshir et al., 2019; Steinwachs et al., 2015). A “pyramidal implementation” of KLT tracker method can be used to detect deformation in the matrix. This method is based on Kanade-Lucas-Tomasi algorithm developed by Kanadé and Lucas which follows bright features from one image to another (Lucas & Kanade, 1981; Tomasi, 1991). It extracts local regions of interest from each image and identifies the corresponding feature in every image. By comparing two consecutive images or an image compared to a reference image, the displacement can be calculated. The pyramid implementation means that first a multi-resolution pyramid of the image intensity and its gradients are computed, before tracking is performed. Then coarse-grained movement is detected on a lower resolution image, and subsequently higher resolution image is taken where the fine movement is detected. The customary code was written for Matlab.

First, the program identifies either bright features or patches with high intensity variations in x and y. Figure 7a shows the detection of bright features in our matrix. The minimum distance as well as the maximum number of points to be detected can be varied. The number of features to detect depends on image size and quality of the matrix signal. Then, the multi-resolution pyramid of the image intensity and its gradients are computed before tracking is performed. The feature detection is performed on a window on the lower resolution image and then on the higher resolution image. After having reached the maximum iteration for all pyramid levels, the displacement of the feature is extracted (between two frames). An overlay of displacement vectors and phase contrast or fluorescent image of cell can be generated (see Figure 7b). This allows to see if the code runs correctly or if artifacts are generated. The calculation can be corrected for drift by using still beads grafted onto the CS as a reference (see section 2.2) or an area where no cell traction force is applied. The drift is then subtracted from the initial mesh displacement result. From then on, further quantification of the matrix displacement can be performed to identify potential patterns.

**Figure 7:**
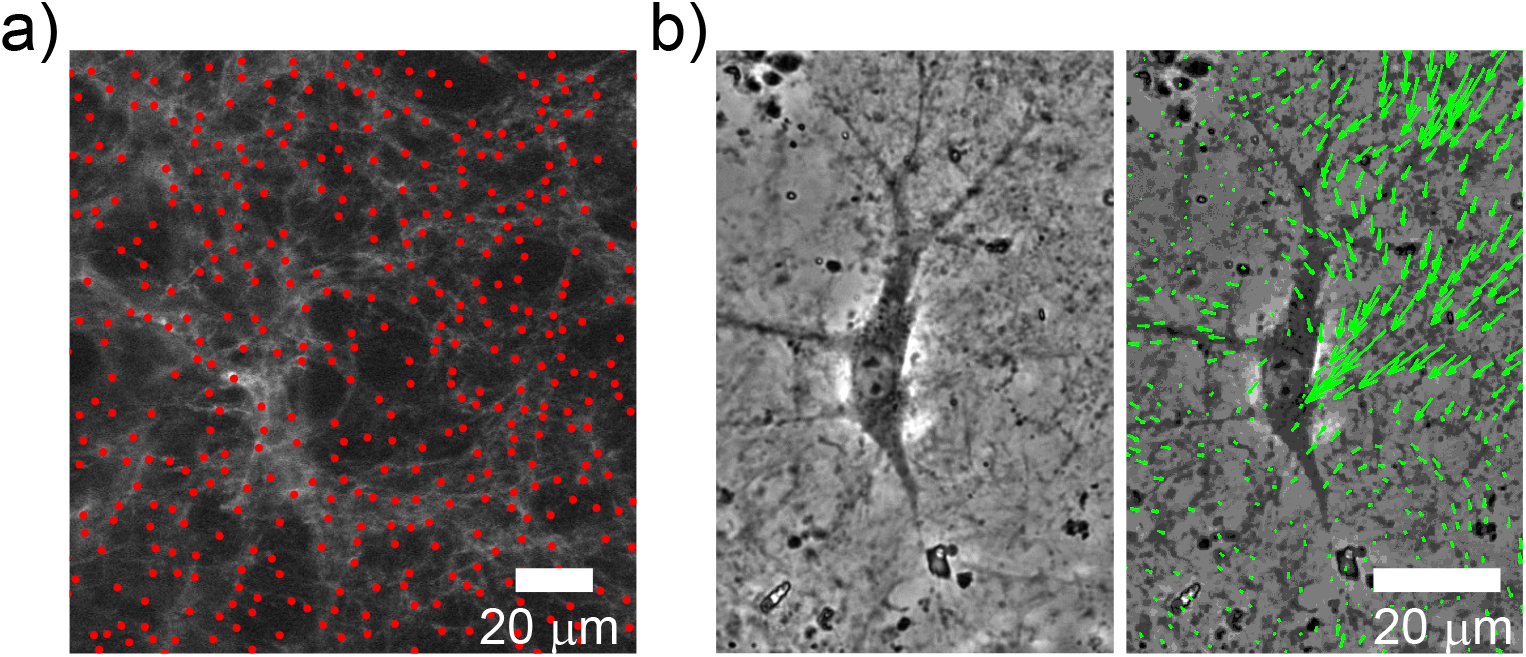
KLT analysis: a) Example of bright point identification of KLT analysis on FN network of CDM with minimum parameter separation of 20 px. b) Snapshot of cell embedded in matrix imaged with phase contrast (left) and the overlay of the matrix displacement (right).

## 4 Discussion

The combination of fluorescent fibronectin within the CDM with the KLT algorithm provide a high accuracy in measurements of displacements for the study of cell motions in 3D physiological environments. Our results indicate that cells apply a specific spatio-temporal force pattern to break the symmetry and migrate (Godeau et al., in preparation). We observed that cells exhibit a specific oscillatory force pattern with localised contraction centers, where a phase shift between front and back contractions accounts for the motion.

In this study, we used NIH 3T3 fibroblasts for CDM culture and for motility assay. In our CDM preparations with NIH3T3 fibroblast, we obtained elastic matrices. Our CDMs were quite soft (~ 50Pa elastic modulus, Godeau et al, in preparation) in comparison to other CDMs reported in the literature (Petrie et al., 2012; Soucy, Werbin, Heinz, Hoh, & Romer, 2011; Tello et al., 2015). Furthermore, we did not detect matrix modification due to cell passage. Our assay may be different for cells which were reported to migrate while deteriorating the ECM (Wolf et al., 2013). These mechanical characteristics are expected to depend on the cell type used for CDM synthesis. Each cell type might incorporate different or a different ratios of extracellular matrix proteins and crosslinkers, and these molecular compositions determine the CDM mechanics.

We tested various cell types which all integrated and migrated within the CDM and deformed the matrix (see Fig. 4). The amplitude of forces applied by cells depend on the cell types. However, all cells integrated in the CDM should go through the spatial network constructed by the CDM-generating cells. This also means that the degree of confinement may vary depending on the cell type. Larger cells in CDMs produced by smaller cells may lead to higher confinement, whereas smaller cells in CDMs produced by larger cells may lead to lower confinement.

## 5 Conclusion

In this chapter, we presented a protocol to generate fluorescent 3D cell derived matrices which allow to perform traction force microscopy and observe key components for the characterization of motion, fibronectin tracking, focal contacts and acto-myosin cytoskeleton. The protocol as it is described was optimised for one cell type (NIH3T3 fibroblasts) but it can easily be adapted to other cell types or combination of cell types (cells producing CDM and cells studied in CDM). We believe that this method could have the potential to address fundamental questions regarding cell movement, cell division and symmetry breaking in physiological environments. It could also be used for diagnostic purposes for comparing cells with different 3D migratory potential and force patterns.

We acknowledge Jordi Comelles, Marco Leoni, Pierre Sens, Sebastien Harlepp, Albrecht Ott, Jacky Goetz, Laszlo Tora and Bernardo Reina San Martin for key insights and help, and we thank the members of Riveline’s lab for experimental support and for stimulating and critical discussions. We thank the Imaging and Cell Culture Platform of IGBMC. This study was supported by the grant ANR-10-LABX-0030-INRT, a French State fund managed by the Agence Nationale de la Recherche under the frame program Investissements d’Avenir ANR-10-IDEX-0002-02. This work was also supported by funds from the CNRS (D.R.), the University of Strasbourg (D.R.), the ci-FRC of Strasbourg (D.R.) and the DFH-UFA (A.G., D.R.).

## List of abbreviations

CDM: Cell Derived Matrix
CS: CoverSlip
ECM: ExtraCellular Matrix
FN: FibroNectin
KLT: Kanade-Lucas-Tomasi

