## Supplementary material for "Generation of fluorescent cell-derived-matrix to study 3D cell migration": Movie Captions

### Movies

Movie 1: Lysis of CDM. Time in mm:ss.

Movie 2: Time-lapse of a NIH3T3 fibroblast migrating in the CDM. Time in hh:mm.

Movie 3: Time-lapse of a HeLa cell migrating in the CDM. Time in hh:mm.

Movie 4: Time-lapse of a mouse lmphocyte migrating in the CDM. Time in hh:mm.

Movie 5: Time-lapse of a mouse embryonic fibroblast migrating in the CDM. Time in hh:mm.

Movie 6: Time-lapse of a REF52 fibroblast migrating in the CDM. Time in hh:mm.

Movie 7: Time-lapse of a NIH3T3 fibroblast entering the CDM. The top shows the bright field image and the bottom the FN mesh. Time in mm:ss.

Movie 8: Time-lapse of a NIH3T3 fibroblast expressing RFP zyxin (red) migrating through the CDM with yellow FN. Time in hh:mm.
